# The geometry of *G* × *E*: how scaling and endogenous treatment effects shape interaction direction

**DOI:** 10.1101/2025.07.15.664999

**Authors:** Michal Sadowski, Andy W. Dahl, Noah Zaitlen, Richard Border

## Abstract

Gene-environment interaction (*G* × *E*) studies hold promise for identifying genetic loci mediating the effects of environmental risk on disease. However, interpretation of *G* × *E* effects is often confounded by two fundamental issues: the dependence of interaction estimates on outcome scale and the presence of endogenous treatment effects, in which genetic liability influences environmental exposure. These factors can induce spurious *G* × *E* signals—even when genetic and environmental contributions are purely additive on an unobserved scale.

In this work, we demonstrate that any monotone convex transformation of an outcome induces *sign-consistent G* × *E* effects: the sign of the interaction term aligns with the sign of the corresponding main genetic effect. We further show that endogenous treatment effects, modeled as threshold-based interventions, generate *G* × *E* effects with a similar directional signature. Exploiting this property, we propose a simple diagnostic: sign consistency across *G* × *E* estimates can identify artifacts driven by outcome scaling or exposure endogeneity.

We validate our framework in the UK Biobank using transcriptome-wide interaction studies (TxEWAS) across multiple trait–environment pairs, observing widespread sign consistency in some settings—suggesting confounding by scaling or treatment bias. Our results provide both a theoretical foundation and a practical tool for interpreting *G* × *E* findings, enabling researchers to distinguish biologically meaningful interactions from those induced by statistical artifacts.

## 1 Introduction

Individuals exhibit substantial phenotypic heterogeneity in response to environmental perturbations. Part of this heterogeneity arises from individual differences in genetic background and is referred to as gene-environment interaction (*G* × *E*). Several interactions identified to date have important implications for human health. For example: (1) dietary treatment prevents symptoms of phenylketonuria—a genetic disorder caused by mutations in the *PAH* gene [1]; (2) physical activity blunts the effects of obesity risk variants in the fat mass and obesity-associated gene, *FTO* [2]; (3) a variant of the *NAT2* gene elevates the risk of bladder cancer in smokers [3]; and (4) a multitude of gene polymorphisms have been shown to impact drug response or toxicity [4–6]. These examples showcase the potential of *G* × *E* discovery to enhance disease prevention and management, to enable design of individualized treatments that are safer and more effective, and, more generally, to advance our understanding of disease etiology. To unlock this potential, many methods for *G* × *E* detection have been developed [7–10] and they are continually being optimized. Most recent approaches enable genome-wide *G* × *E* screens in large-scale studies of human populations [11–13].

However, two fundamental issues complicate the interpretation of current *G* × *E* approaches: (1) dependence on phenotype scale and (2) endogenous treatment effects. In case (1), the detection of an interaction effect and its direction depend on the scale on which the outcome is measured, or to which it might be transformed [14–17]. For example, even though a genetic variant *G* and an environmental factor *E* impact an outcome *Y* additively, an interaction test performed on *Y* that has been log-transformed (e.g. as part of quality control processing) can yield a highly significant *G* × *E* effect (Figure 1). More generally, many interaction effects can be induced or removed by monotonic non-linear transformations of the data. In case (2), exposure and genetic liability are causally intertwined. For example, imagine that a treatment is administered to taper the level of a heritable phenotype when it crosses some threshold (e.g. statins may be prescribed to lower low-density lipoprotein (LDL) cholesterol levels). In this case, exposure to the intervention is related to genetic factors influencing phenotype. Such endogenous treatment effects can result in apparent *G* × *E*, even when gene and environment act additively on the observed scale. As a result, most *G* × *E* findings require the caveat that they may be a consequence of measurement scale and/or endogenous treatment effects [18–21].

**Figure 1:**
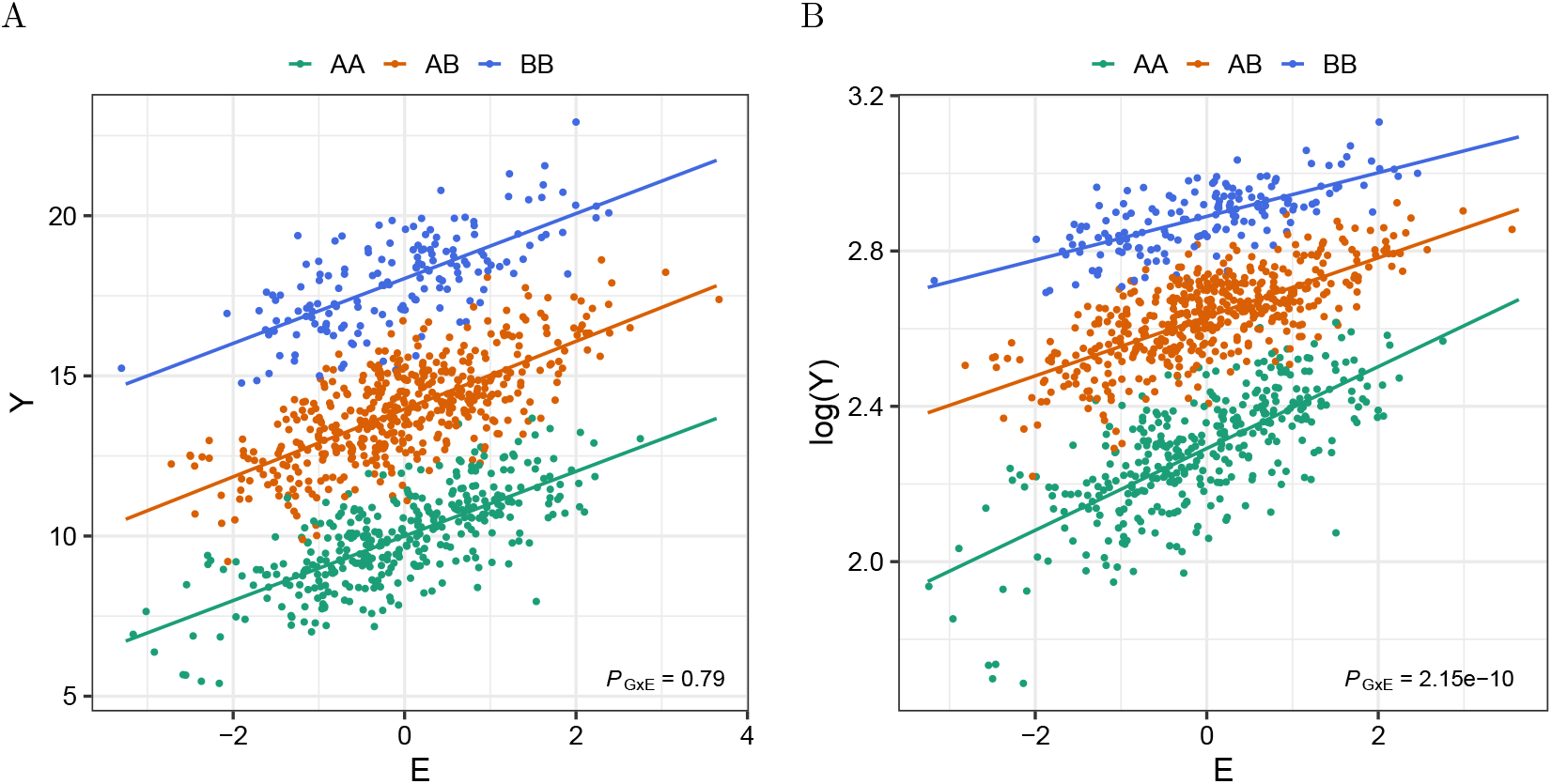
*G* × *E* effect induced by log-transformation of outcome *Y*. (A) Depiction of the effects of the genetic variant *G* (with reference allele A and alternative allele B, MAF=0.4), the environmental factor E (drawn from a standard normal distribution), and the interaction between the two (*G* × *E*) on the outcome *Y* (generated as *Y* ∼ 𝒩(10 + 4*G* + *E*, 1) for 1,000 samples). The p-value for the *G* × *E* effect (*P*_G×E_) is given in the right lower corner. (B) Depiction of the same effects on the log-transformed *Y*. Whereas the *G* × *E* effect is not detectable for *Y* (A), it is detectable for the log-transformed *Y* (B).

Here, we demonstrate that monotone convex transformations of an outcome always induce *sign-consistent G* × *E*, where the direction of the interaction effects is determined by the sign of the corresponding main effects. We further show that a simple model of endogenous treatment effects also generates sign-consistent *G* × *E*. Finally, we discuss examples of non-convex transformations, like the logistic function, showing why and under what circumstances they induce this particular type of interaction effect. Our results indicate that a simple examination of sign consistency across detected *G* × *E* can rule out the possibility that all interactions have been induced by a monotone convex scaling of the outcome or endogenous treatment effects. Another consequence of this result is that if the *G* × *E* signal is not only a scaling artifact, only a fraction of this signal can be eliminated by a single scaling of the data. We demonstrate the usefulness of sign consistency examination in real data, as our analysis of this property identifies that statin use induces false positive gene-age interaction effects on LDL cholesterol levels.

## 2 Sign-consistent interaction property

We investigate the relationship between the signs of regression coefficients estimated in the *G* × *E* model across multiple genetic variants. More concretely, consider testing two haploid variants, *G*_1_ and *G*_2_, for an interaction with a binary environmental exposure *E* against phenotype *Y* in two single-variant regressions:

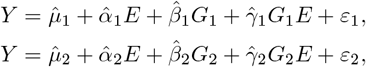

where 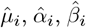 and 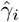 are coefficients estimated in regression *i*, and *ε*_*i*_ represents residual variation. Consider also the same two tests performed for the same phenotype *Y* measured on a different scale:

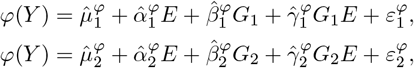

where *φ* is the map from the scale of the former measurement of *Y* to the scale of the latter measurement of *Y*, and the corresponding coefficients and residuals are marked with superscript *φ*.

We demonstrate that the signs of interaction effects, 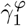 and 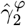, induced by a monotone convex transformation *φ* depend on the signs of the main genetic effects, 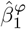 and 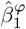. Formally:

### Claim 1

(Sign-consistent interaction property). *Assume measurement Y has homogeneous variance and exhibits no G* × *E effects* 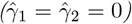 *on the original scale, and function φ is monotone convex, then the G* × *E effects* 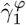 *and* 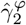 *are consistently oriented with respect to the corresponding main genetic effects:*

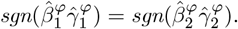

Note that if we assume that the alleles of *G*_1_ and *G*_2_ are encoded so that their main effects have the same direction (i.e., 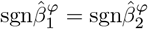), the above property becomes:

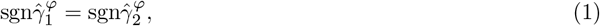

which we call *sign consistency*. By contraposition, if the homoskedasticity and 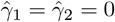 assumptions hold but property (1) does not, non-zero *G* × *E* effects 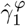 and 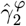 are not both induced by a monotone convex scaling of *Y*.

Generally, in abstraction from this two-variant example, we demonstrate that if there is a scale, on which phenotype *Y* has homogeneous variance across values of environmental factor *E* and genetic variant *G*, and those factors have only additive effects on *Y*, then the direction of the *G* × *E* effect estimated for a measurement that is a monotone convex transformation of *Y* depends on the direction of the corresponding main genetic effect.

We, therefore, propose to examine observed *G* × *E* effects for the sign consistency property, as it provides a means to exclude the family of monotone convex transformations as the sole source of these effects.

In what follows, we first outline the proof of the presented property for the family of monotone convex transformations; the full proof is given in Appendix A. We then demonstrate that a simple model of endogenous treatment effects also generates consistently oriented *G* × *E* and can be evaluated using the same property. We next discuss why the presented property need not hold for non-convex transformations in general. Finally, we examine sign consistency of *G* × *E* effects in real data, and end with a discussion of the implications of our theoretical results for interpretation of *G* × *E* studies.

## 3 Sketch of argument

Suppose that phenotype *Y* has homogeneous variance across values of environmental factor *E* and genotype *G*:

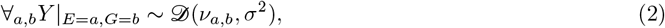

where 𝒟 is a symmetric distribution with mean *ν* and variance *σ*^2^. We fit the following linear regression model to test the effect of an interaction between *G* and *E* on this phenotype:

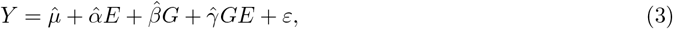

where 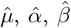 and 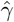 are estimated coefficients and *ε* is the error. We consider a simplified example, where *E* and *G* are binary variables, which corresponds to the case of a haploid genetic variant and a binary environmental exposure. Importantly, our reasoning is not contingent on whether this model is correct, only that the homoskedasticity condition (2) holds. The coefficients in (3) can be related to the empirical conditional expectations:

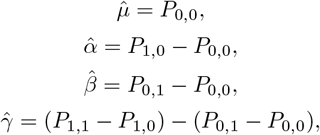

Where 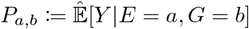. For example, *P*_0,1_ is the average value of phenotype *Y* in individuals with genotype *G* = 1 who are not exposed to environmental factor *E* (i.e., *E* = 0).

Without loss of generality, suppose further that coefficients 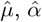 and 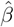 are estimated to be positive, and that the estimate of the *G* × *E* effect 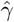 is zero, as depicted on the x-axis of Figure 2A. Consider now a regression similar to (3), but performed on phenotype *Y* measured on a different than the original scale:

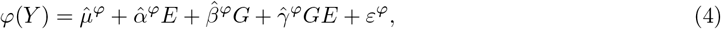

where *φ* is the map from the original scale of *Y* to the new scale.

**Figure 2:**
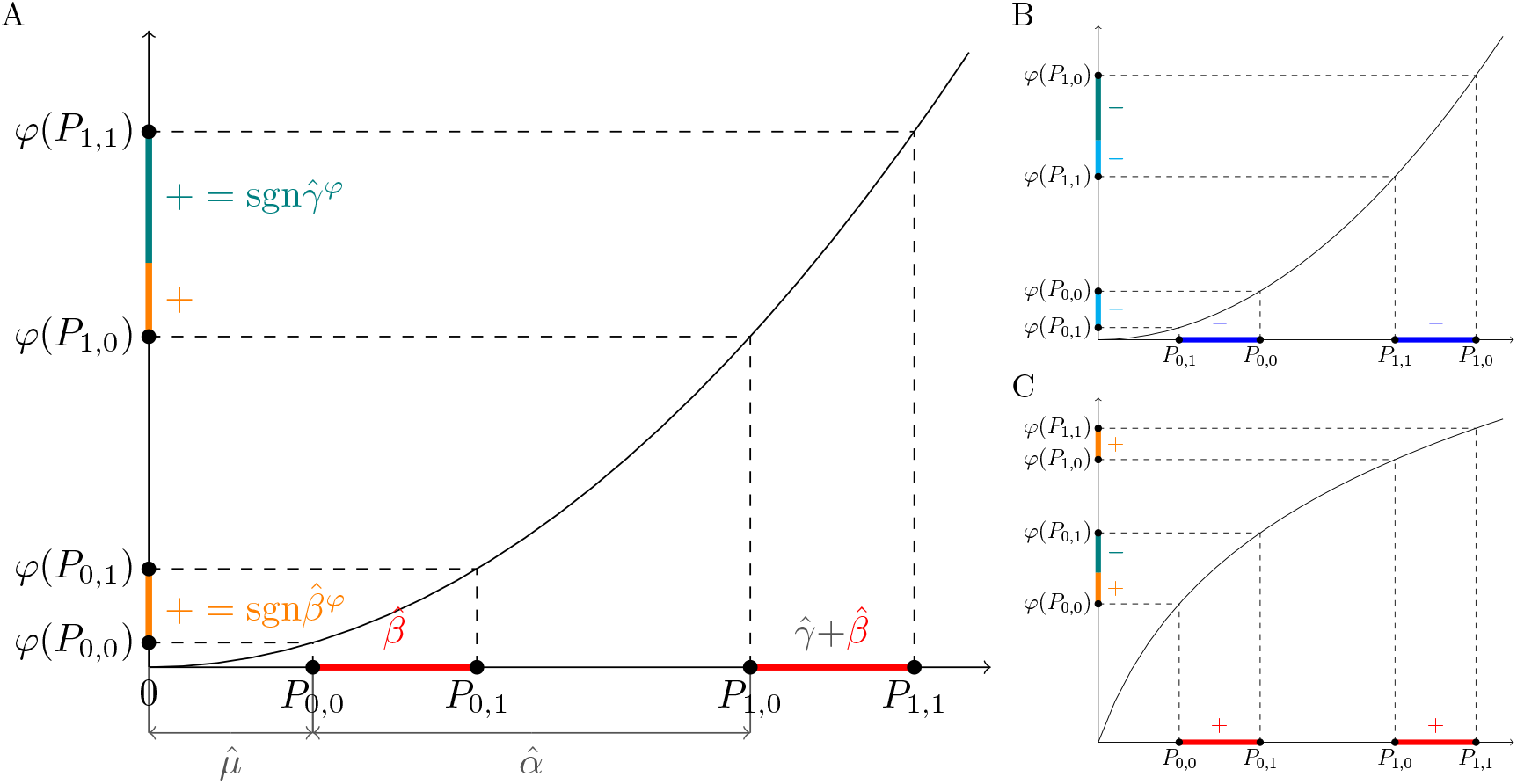
Monotone convex transformations of the outcome induce sign-consistent *G* × *E* effects in interaction tests. (A) The x-axis shows the intercept 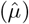, the main effect of 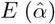, the main effect of *G* 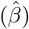 and the *G* × *E* effect 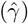 estimated by regressing *E, G* and *GE* on phenotype *Y*. If, as assumed here, 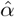 and 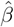 are positive and 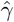 is null, a similar regression on this phenotype transformed with an increasing convex down function *φ* will yield the main effect of 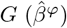 and the *G* × *E* effect 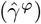 that are positive. The sign of the *G* × *E* effect can be calculated by 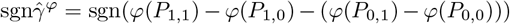, where 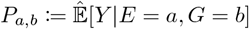. (B) Similar to (A), but shows the signs of 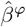 and 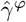 when 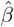 is negative. (C) Similar to (A), but shows the signs of 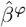 and 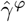 when *φ* is increasing convex up.

If we assume that *φ* is increasing convex down (Figure 2A), then the signs of 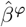 and 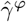 can be related to points *P*_*a,b*_ as:

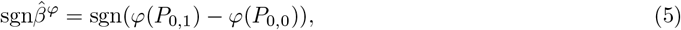

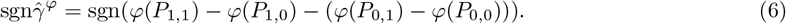

That is, with the above assumptions, the direction of the effects estimated for the scaled phenotype *φ*(*Y*) can be expressed using quantities *P*_*a,b*_ defined on the original scale of *Y* (see Appendix A for a derivation of this fact). Looking at Figure 2A, it is easy to see that in our example:

1. The signs of differences *φ*(*P*_0,1_) − *φ*(*P*_0,0_) and *φ*(*P*_1,1_) − *φ*(*P*_1,0_) are the same, and, by (5), follow the sign of 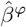.
2. The magnitude of *φ*(*P*_1,1_) − *φ*(*P*_1,0_) is larger than the magnitude of *φ*(*P*_0,1_) − *φ*(*P*_0,0_).

From (6) and these two facts it follows that the sign of 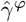 is positive. Note that this will be true for any genetic variant whose main effect, 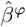, tested in (4) is positive. If, on the other hand, 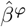 is negative, the *G* × *E* effect, 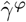, will be negative (Figure 2B). In general, we have the following relation:

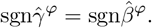

Applying a similar argument, it can be shown that when *φ* is increasing convex up, the opposite relation, 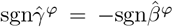, holds (Figure 2C). In general, the direction of this relation depends on whether *φ* is increasing or decreasing and convex down or convex up, and on the sign of 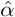, that is, the estimated effect of environmental factor *E* on phenotype *Y* (Table 1). A complete proof discussing all these cases is given in Appendix A.

**Table 1:** Increasing or decreasing and convex down or convex up transformations induce *G* × *E* effects 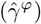 whose directions are consistent with the directions of the observed main genetic effects 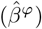.

| | $\hat{\alpha} > 0$ | $\hat{\alpha} < 0$ |
| --- | --- | --- |
| $\varphi$ increasing convex down | $\text{sgn}\hat{\gamma}^\varphi = \text{sgn}\hat{\beta}^\varphi$ | $\text{sgn}\hat{\gamma}^\varphi = -\text{sgn}\hat{\beta}^\varphi$ |
| $\varphi$ decreasing convex down | $\text{sgn}\hat{\gamma}^\varphi = -\text{sgn}\hat{\beta}^\varphi$ | $\text{sgn}\hat{\gamma}^\varphi = \text{sgn}\hat{\beta}^\varphi$ |
| $\varphi$ increasing convex up | $\text{sgn}\hat{\gamma}^\varphi = -\text{sgn}\hat{\beta}^\varphi$ | $\text{sgn}\hat{\gamma}^\varphi = \text{sgn}\hat{\beta}^\varphi$ |
| $\varphi$ decreasing convex up | $\text{sgn}\hat{\gamma}^\varphi = \text{sgn}\hat{\beta}^\varphi$ | $\text{sgn}\hat{\gamma}^\varphi = -\text{sgn}\hat{\beta}^\varphi$ |

## 4 Endogenous treatment effects

Suppose that the environmental factor *E* tested in the *G* × *E* regression model is a treatment. Suppose that this treatment is administered to taper the level of a heritable phenotype when it crosses some threshold—e.g. the statin therapy for individuals with high LDL cholesterol. In this case, exposure to the intervention is related to genetic factors influencing phenotype—which we refer to as *endogenous treatment effects*. As shown in [6], endogenous treatment effects can cause false discoveries when the observed levels of the phenotype (subjected to treatment) are tested for *G* × *E*. Following, we prove that the *G* × *E* effects induced in a simple model of endogenous treatment effects are sign-consistent.

Consider the following model of phenotype *Y* :

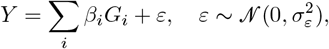

where, again, *G*_*i*_ indicates the presence of an alternative allele at haploid variant *i, β*_*i*_ is the effect of this allele on *Y*, and *ε* is the environmental noise. We assume that the environmental noise is homoskedastic (*∀*_*i*_Var[*ε*|*G*_*i*_ = 0] = Var[*ε*|*G*_*i*_ = 1]).

Suppose that if the level of *Y* is high, an individual is administered treatment *E*:

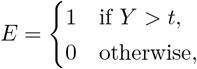

where *t* is some threshold. When applied, treatment *E* changes the level of *Y* by *α*:

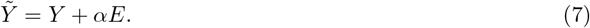

### Claim 2

(Endogenous treatment effect interaction sign property). *Suppose that we observe phenotype* 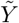 *and test the effect* 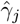 *of the interaction between variant G*_*j*_ *and environmental factor E on this phenotype:*

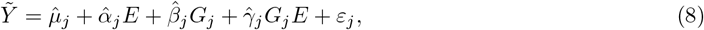

*Then the direction of the estimated G* × *E effect*, 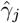, *is opposite to the direction of the main genetic effect*, 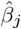.

As in Section 3, we define quantities *P*_0_ and *P*_1_ on the scale of Y, which, transformed, can be used to compute the main genetic effect, 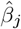, and the *G* × *E* effect, 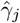, on the scale of 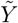. Note that in this case, not only the signs, but also the values of these effects can be expressed in terms of *P*_0_ and *P*_1_ transformed by a certain function. We investigate the properties of this function to determine the properties of 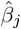 and 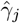. Specifically, we define *P*_0_ ≔ *t* − 𝔼[*Y* |*G*_*j*_ = 0] and *P*_1_ ≔ *t* − 𝔼[*Y* |*G*_*j*_ = 1], and show (Appendix B) that the main genetic effect 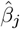 and the *G* × *E* effect 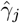 estimated in (8) can be related to these points as:

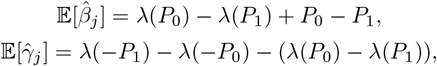

where *λ* is the inverse Mill’s Ratio (*λ*(*x*) ≔ *ϕ*(*x*)*/*Φ(*x*), where *ϕ* and Φ are the standard normal probability density function and cumulative distribution function, respectively). Importantly, *λ* is decreasing and strictly convex down [22].

Suppose that 0 *< P*_0_ *< P*_1_. Then, looking at Figure 3, we see that:

1. The difference *λ*(*P*_0_) − *λ*(*P*_1_) has the opposite sign to difference *P*_0_ − *P*_1_.
2. The magnitude of *P*_0_ − *P*_1_ is larger than the magnitude of *λ*(*P*_0_) − *λ*(*P*_1_).
3. The difference *λ*(−*P*_1_) − *λ*(−*P*_0_) has the same sign as difference *λ*(*P*_0_) − *λ*(*P*_1_).
4. The magnitude of *λ*(−*P*_1_) − *λ*(−*P*_0_) is greater than the magnitude of *λ*(*P*_0_) − *λ*(*P*_1_).

**Figure 3.**
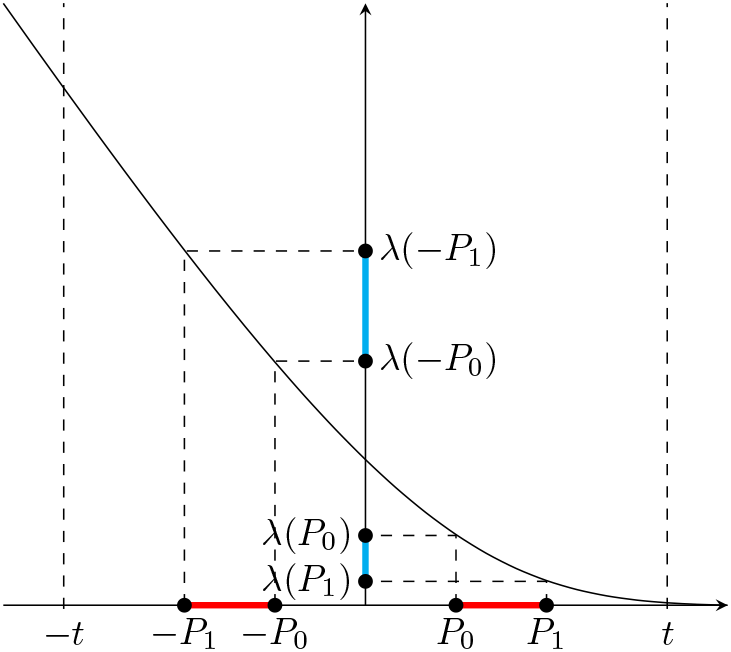
Endogenous treatment effects induce sign-consistent *G* × *E* effects. The x-axis shows the quantities *P*_0_ ≔ *t* − 𝔼[*Y* |*G* = 0] and *P*_1_ ≔ *t* − 𝔼[*Y* |*G* = 1] defined for phenotype *Y* affected by the haploid genetic variant *G*. The main effect of *G* and the effect of *GE* on phenotype *Y*, after treatment *E* is applied to reduce levels of *Y* that exceed threshold *t*, can be expressed as functions of *P*_0_ and *P*_1_ and their images under the inverse Mill’s Ratio *λ*(*P*_0_) and *λ*(*P*_1_). The signs of those functions are dependent.

From our assumption that 0 *< P*_0_ *< P*_1_ and facts 1 and 2 above, it follows that the sign of 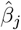 is negative. From the same assumption and facts 1, 3 and 4, it follows that the sign of 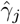 is positive. Note that once the sign of difference *P*_0_ − *P*_1_ is established, the signs of 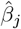 and 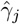 can be determined based on the properties of *λ*. In our example, *P*_0_ − *P*_1_ is positive, which makes 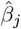 negative and 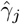 positive. On the other hand, when *P*_0_ − *P*_1_ is negative, 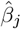 is positive and 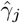 is negative. Therefore, we have shown that *G* × *E* effects induced by endogenous treatment effects (modeled as in (7)) have opposite directions to the corresponding main genetic effects. That is, for any *j*, 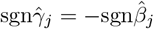.

The same property holds when treatment *E* is administered whenever the level of *Y* is below threshold *t* (Appendix B).

## 5 Non-convex transformations

An arbitrary non-linear scaling of an outcome may or may not induce sign-consistent *G* × *E*. To provide more intuition on this, we discuss properties of *G* × *E* that can be produced by two examples of non-convex scaling commonly used in genetic analyses: (1) the logistic function and (2) the inverse normal transformation (INT).

Specifically, we are interested in the relationship between the signs of the main effect 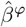 of genetic variant *G* and the effect 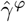 of the interaction between *G* and environmental factor *E* on *φ*-transformed phenotype *Y* in the linear regression:

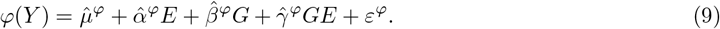

Following Section 3, we assume that *E* and *G* are binary and that *Y* (on the original scale) has homogeneous variance and does not exhibit *G* × *E* effects, meaning that the linear regression:

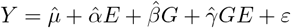

yields 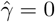.

As demonstrated earlier, given these assumptions, 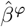 and 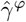 can be defined in terms of the empirical conditional means of untransformed *Y* :

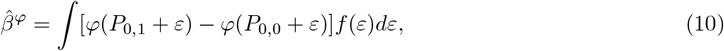

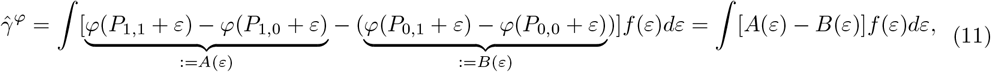

where 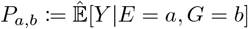 and *f* is the PDF of *ε*.

### Case study: the logistic function

Let *φ* be a logistic function:

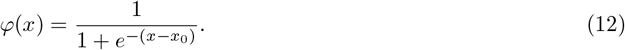

Using definitions (10) and (11), it is easy to see that the sign of 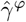 induced by the logistic scaling depends not only on the relative positions of points *P*_0,0_ and *P*_1,0_ (as was the case for monotone convex transformations), but also on their values and the width of the distribution of *ε* (Figure 4A). More specifically, the values of *P*_0,0_ and *P*_1,0_ determine—up to noise—the relation between the magnitudes of *A*(*ε*) and *B*(*ε*) in (11). Since these values will be different for different genetic variants, the relationship between the signs of 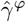 and 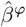 can be variant-specific.

**Figure 4:**
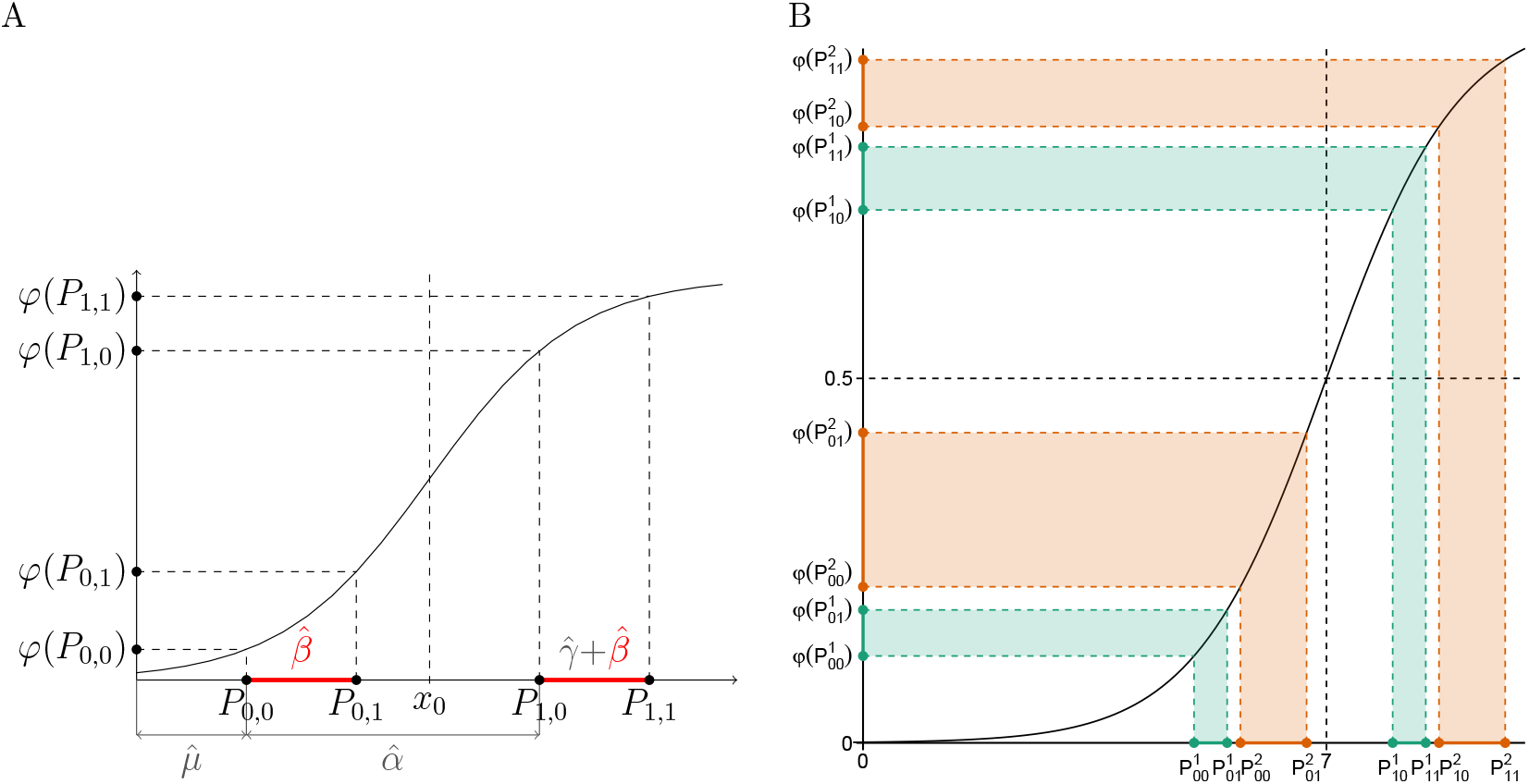
Logistic transformation of the outcome induces *G* × *E* effects that may or may not be sign-consistent. (A) Depiction of the effect that the logistic transformation 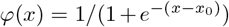 of the outcome may have on the regression-based *G* × *E* test. Compare with Figure 2. (B) An example of two genetic variants (green and orange) with positive effects on the phenotype that after transforming this phenotype with the logistic function *φ*(*x*) = 1*/*(1 + *e*^−(*x*−7)^) exhibit *G* × *E* effects of opposite directions. Compare with Figure 2.

Consider an illustrative example of two genetic variants *G*_1_ and *G*_2_. For simplicity, we assume *ε* = 0, which simplifies (10) and (11) to:

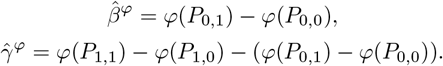

For *G*_1_ we assume: 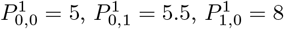 and 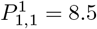. This means that in unexposed individuals carrying the reference allele at *G*_1_ the average value of phenotype *Y* is 5, whereas in exposed individuals carrying the same allele it is 8; and the effect of *G*_1_ in both unexposed and exposed groups is 0.5. For variant *G*_2_, on the other hand, we assume: 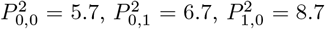 and 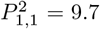, which corresponds to average values of 5.7 and 8.7 in unexposed and exposed non-carriers, respectively, and the genetic effect of 1 (see x-axis of Figure 4B). Now imagine that we convert phenotype *Y* to a risk scale where the value of 7 corresponds to the risk of 50%: *φ*(*Y*) = 1*/*(1 + *e*^−(*Y* −7)^), and perform regression (9). For variant *G*_1_, this regression yields a positive main genetic effect and a positive *G* × *E* effect. For variant *G*_2_, it produces a positive main effect, but a negative *G* × *E* effect (Figure 4B). Thus, in this example the logistic transformation induces *G* × *E* effects that are not sign-consistent.

There are, however, scenarios where the logistic scaling will induce *G* × *E* that are sign-consistent. A plausible example of such a scenario in healthcare data occurs when *x*_0_ in (12) is large (meaning that the cases are called at high phenotype values) and the environmental effect and individual genetic effects on (untransformed) *Y* are relatively small, so that all points *P*_*a,b*_ for all considered genetic variants are smaller than *x*_0_. Since the logistic function is convex on the domain (− inf, *x*_0_], transforming points *P*_*a,b*_ with this function yields a relation 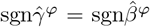 (Table 1). In general, the sign of a *G* × *E* effect induced by the logistic scaling depends on the relative positions of *P*_0,0_, *P*_1,0_, *P*_0,1_ and *P*_1,1_ with respect to *x*_0_—all possible cases are detailed in the Supplementary Material.

### Case study: the inverse normal transformation

Another data transformation commonly used in genetic analyses is the INT. It matches quantiles of the data distribution with the quantiles of the standard normal distribution; thus it preserves the order of data points, but not the distances between them. In particular, the distances between conditional means *P*_0,0_, *P*_0,1_, *P*_1,0_ and *P*_1,1_ transformed by INT will depend on their values, which are different for different variants tested in (9). As a consequence, the INT transformation can induce *G* × *E* effects in any direction with respect to the main genetic effect of a given sign.

## 6 Previously published interaction results exhibit sign consistency property

We have examined sign consistency for several *G* × *E* studies, selecting *E*-outcome pairs for which interactions have previously been found (Figure 5A). More specifically, we performed TxEWAS [6, 23] in the UK Biobank [24] population of unrelated white British individuals (Supplementary Material). TxEWAS tests the effect of the interaction between predicted expression of a gene *G* and environmental exposure *E* on phenotype *Y* using the following linear regression model:

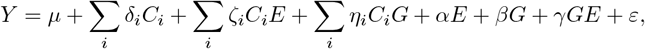

where *C*_*i*_ is the *i*-th additional environmental covariate included in the model, greek letters represent effect sizes, and *ε ∼* 𝒩 (0, *σ*^2^). Among additional covariates we included: age, sex, birth date, Townsend deprivation index, and the first 16 genetic principal components (PCs) [25] (if not already used as *E*). In a single study, we performed multiple tests for a single gene—corresponding to multiple tissues in which this gene was expressed—and used the hierarchical FDR (hFDR) correction to call significant interactions from aggregated results [6]. Sign consistency was examined considering these interactions in tissues, in which they had the strongest effects.

**Figure 5:**
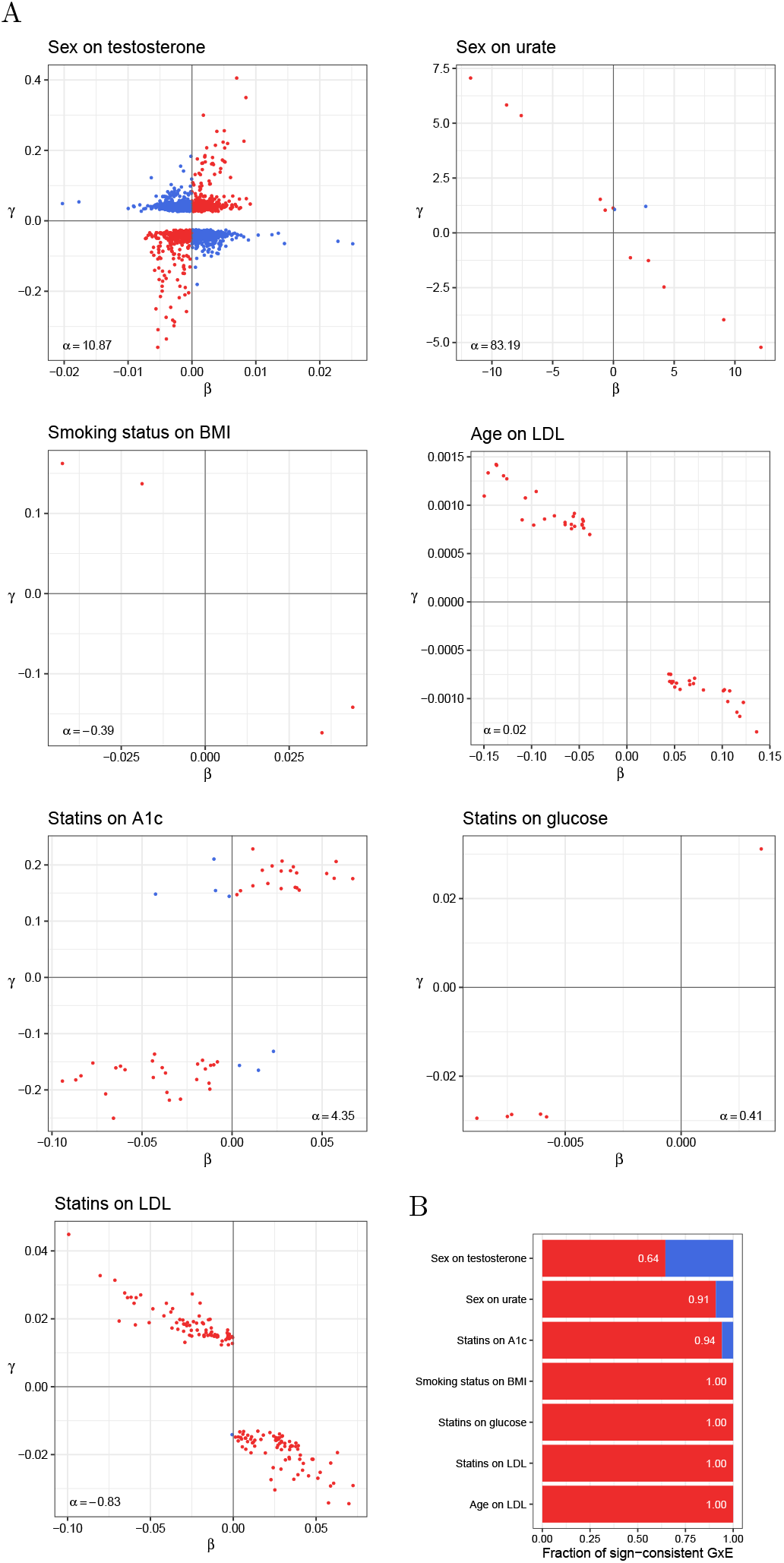
Sign consistency between the main (*β*) and interaction (*γ*) effects in TxEWAS for select *E*’s and outcomes. (A) Main vs interaction effects for identified genes. For each gene, we plot the estimates corresponding to the tissue with the strongest interaction p-value. *α* is the main environmental effect. (B) The fraction of *G* × *E* that have sign-consistent effects. This fraction was calculated among interacting genes (called at hFDR *<* 10%) whose main effects were nominally significant at 5%. The tissue with the strongest interaction p-value for a given gene was considered.

We have investigated gene-sex interaction effects on the primary male sex hormone, testosterone, and the end-product of the purine metabolism, urate; gene-smoking interaction effects on body mass index (BMI); gene-age interaction effects on LDL cholesterol levels; and gene-statins interaction effects on statins’ primary target, LDL cholesterol, and phenotypes related to their potential side effect on diabetes risk [26, 27]—blood glucose and hemoglobin A1c (see Supplementary Material for phenotype definitions and preprocessing details). We have observed moderate to strong evidence for sign-consistent *G* × *E* effects across these traits. More specifically, the fraction of sign-consistent *G* × *E* effects was moderate for the sex-testosterone *E*-outcome pair, high for the sex-urate and statins-hemoglobin A1c pairs, and maximal for the rest of our studies (Figure 5B).

This suggests the interaction effects identified in most of these studies may have been induced by the outcome measurement scaling or endogeneity. For example, we hypothesized that the interaction effects detected in the age-LDL cholesterol study were a consequence of endogenous treatment effects. This is because with age increases the probability of taking statins, which are prescribed at high LDL cholesterol levels—meaning that genetic variation associated with LDL cholesterol levels is also correlated with age. Indeed, when we included statin use in our model as a covariate, the *G* × *E* effects disappeared. This is not to say that all *G* × *E* effects identified in studies that exhibited perfect sign consistency should be presumed artifacts. For example, many gene-statins effects identified by TxEWAS for LDL cholesterol were replicated in a retrospective longitudinal pharmacogenomic study [6]. Nonetheless, this analysis demonstrates that examination of sign consistency can be useful in explaining observed statistical interactions.

## 7 Discussion

We have demonstrated that if there is a scale on which an outcome has homogeneous variance across values of environmental factor *E* and genetic variant *G*, and these factors have only additive effects on this outcome, then the direction of the *G* × *E* effect estimated on the scale that is a monotone convex transformation of the original outcome scale is determined by the direction of the main effect of *G*. In addition, we have shown that endogenous treatment effects, modeled as threshold-based interventions, can only produce *G* × *E* effects with the same sign property.

A consequence of our result is that if *G* × *E* effects in both directions with respect to the main genetic effects are observed, there is no monotone convex transformation that can eliminate the *G* × *E* effects. Furthermore, they could not have been all induced by endogenous treatment effects. Our results are related to prior conditions under which outcome scaling can eliminate interaction effects [16], especially prior results bounding interaction effect sizes as a function of the curvature of the scaling function [28].

Exploiting this property, we propose that researchers examine sign consistency across observed *G* × *E* effects to assess the possibility that interactions are driven by monotone convex scaling of the outcome or endogenous treatment effects. Our analysis of real data sets demonstrates that such an examination can help detect possible confounding. It can also be useful for ruling out the possibility that identified interactions are scaling or endogeneity artifacts.

In principle, the proportion of sign-consistent interaction effects observed in a *G* × *E* study may be underestimated due to the noise in the estimates. In our analysis such inaccuracy, if present, is small since we consider statistically significant effects only.

We note that the homoskedasticity assumption made in our proofs is also an assumption of the linear regression model. Violation of this assumption results in a biased test for the interaction effect [6, 29]. In the observed data, it is specifically common that the variance of the outcome differs across strata defined by the environmental factor [30]. Owing to its importance and incomplete characterization, we comprehensively examine the conditional heteroskedasticity bias in the Supplementary Material. We analytically describe the conditions under which this bias is expected to arise and the direction of its effect. It has been established that, in the presence of heteroskedasticity, *G* × *E* should be modeled using the double generalized linear model or a standard linear model modified to incorporate robust standard errors [6, 29].

## Appendix A

Suppose that there is a scale, on which a phenotype exhibits no *G* × *E*, and has homogeneous variance. We show that any monotone convex transformation of this phenotype can only induce sign-consistent *G* × *E* effects (Claim 1). The implication is that if *G* × *E* effects in both directions with respect to the main genetic effects are observed, there is no such transformation that can eliminate the *G* × *E* effects.

Specifically, consider phenotype *Y* that has homogeneous variance across values of binary environmental factor *E* and haploid genotype *G*:

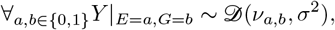

where 𝒟 is a symmetric distribution with mean *ν* and variance *σ*^2^. Consider further fitting the following linear regression model to *Y* :

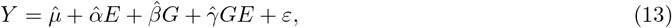

where 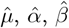 and 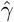 are estimated coefficients, and *ε* is the error. We assume that:

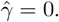

The coefficients in (13) can be related to the empirical conditional means of *Y*, which we denote by points 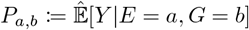:

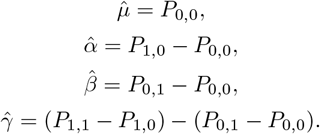

The order of *P*_0,0_ and *P*_0,1_, *P*_1,0_ and *P*_1,1_, *P*_0,0_ and *P*_1,0_, and *P*_0,1_ and *P*_1,1_ is determined by the signs of coefficients 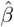 and 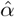. To see this, note that if 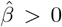, then *P*_0,0_ *< P*_0,1_. Alternatively, if 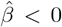, then *P*_0,0_ *> P*_0,1_. Furthermore, by the assumption that 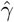 is null, 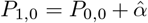 and 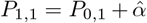 (Figure 6).

**Figure 6:**
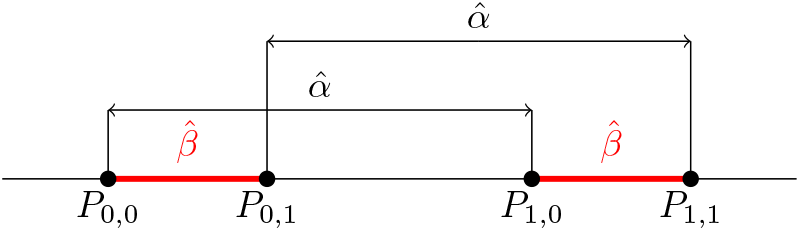
When 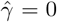, the signs of regression coefficients 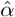 and 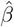 in (13) determine the order of *P*_0,0_ and *P*_0,1_, *P*_1,0_ and *P*_1,1_, *P*_0,0_ and *P*_1,0_, and *P*_0,1_ and *P*_1,1_. Here, 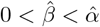.

We will use this fact to show that a regression similar to (13) on a monotone convex transformation of *Y* yields *G* × *E* effects whose directions depend on the directions of the corresponding main genetic effects.

Consider *Y* transformed by a function *φ*, and a linear regression of this transformed *Y* on *E, G* and *GE*:

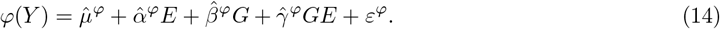

We can relate the coefficients 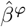 in (14) to points *P*_0,0_, *P*_0,1_, *P*_1,0_, and *P*_1,1_ that we have defined on the original scale of *Y* :

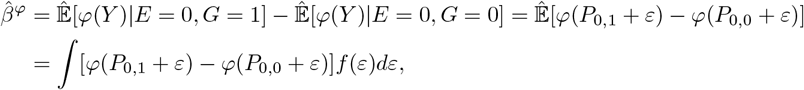

and likewise for 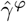:

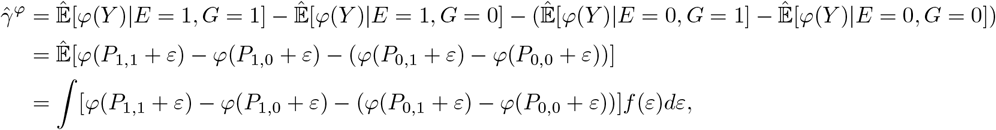

where *f* is the PDF of *ε*. Note that each point *P*_*a,b*_ above is always shifted by the same value; and that the signs of the above expressions are invariant to this shift if *φ* is monotone convex:

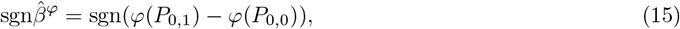

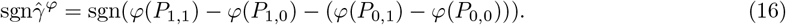

Without loss of generality, suppose that *φ* is increasing convex down. To determine the sign of 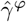, we need to know the signs of differences *φ*(*P*_0,1_) − *φ*(*P*_0,0_) and *φ*(*P*_1,1_) − *φ*(*P*_1,0_), and the relation between their magnitudes. Since *P*_0,1_ and *P*_1,1_ are shifted from *P*_0,0_ and *P*_0,1_ by the same value, 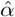, the signs of *φ*(*P*_0,1_) − *φ*(*P*_0,0_) and *φ*(*P*_1,1_) − *φ*(*P*_1,0_) are the same, and, by (15), follow the sign of 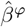 (Figure 7A). The relation between their magnitudes depends on the sign of 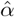. If 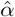 is positive, the magnitude of *φ*(*P*_1,1_) − *φ*(*P*_1,0_) is greater than the magnitude of *φ*(*P*_0,1_) − *φ*(*P*_0,0_), and the opposite is true if 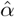 is negative.

**Figure 7:**
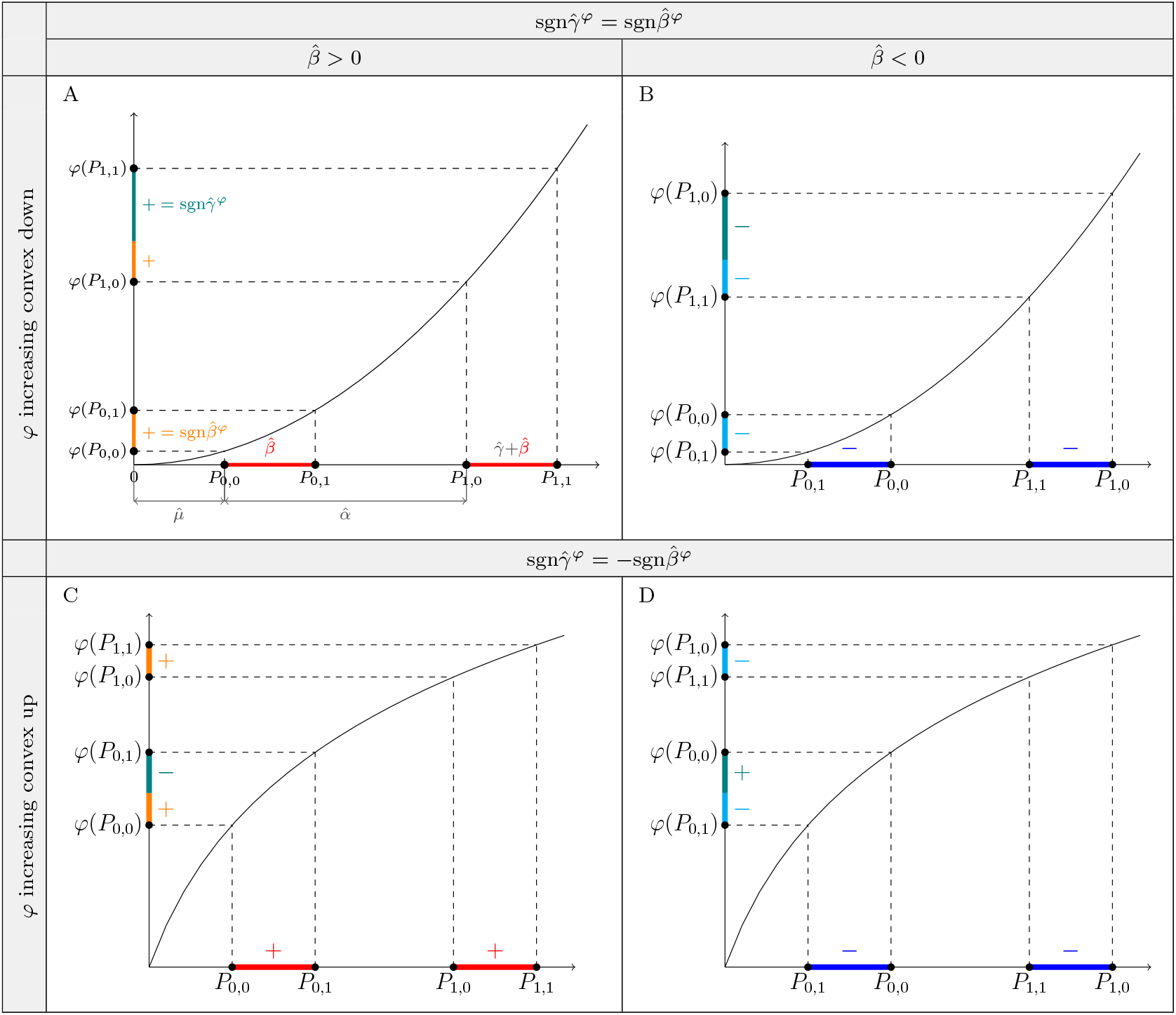
Increasing convex transformations of the outcome induce *G* × *E* effects, 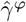, whose direction is determined by the direction of the main genetic effects, 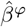, if the untransformed outcome exhibits no *G* × *E* and has homogeneous variance in regression (13). (A) The relation between the signs of 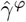 and 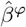 when 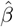 and 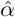 are positive and *φ* is increasing convex down. (B) Similar to (A), but when 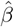 is negative. (C) Similar to (A), but when *φ* is increasing convex up. (D) Similar to (A), but when 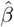 is negative and *φ* is increasing convex up.

If two genetic variants, *G*_1_ and *G*_2_, are regressed like *G* in (14)—and the assumptions of model (13) are met—such that the sign of 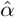 in these two cases is the same, the sign of 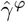 differs between these regressions only if the sign of 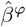 differs.

For example, when *φ* is increasing convex down and 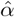 is positive, then:

- 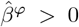 implies 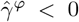, because both *φ*(*P*_0,1_) − *φ*(*P*_0,0_) and *φ*(*P*_1,1_) − *φ*(*P*_1,0_) are positive, and |*φ*(*P*_0,1_) − *φ*(*P*_0,0_)| *<* |*φ*(*P*_1,1_) − *φ*(*P*_1,0_)| (Figure 7A).
- 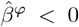 implies 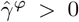, because both *φ*(*P*_0,1_) − *φ*(*P*_0,0_) and *φ*(*P*_1,1_) − *φ*(*P*_1,0_) are negative, and |*φ*(*P*_0,1_) − *φ*(*P*_0,0_)| *<* |*φ*(*P*_1,1_) − *φ*(*P*_1,0_)| (Figure 7B).

Therefore, in this example, 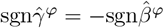.

Similarly, when 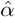 is positive, but *φ* is increasing convex up, then:

- 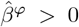 implies 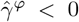, because both *φ*(*P*_0,1_) − *φ*(*P*_0,0_) and *φ*(*P*_1,1_) − *φ*(*P*_1,0_) are positive, and |*φ*(*P*_0,1_) − *φ*(*P*_0,0_)| *>* |*φ*(*P*_1,1_) − *φ*(*P*_1,0_)| (Figure 7C).
- 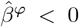 implies 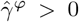, because both *φ*(*P*_0,1_) − *φ*(*P*_0,0_) and *φ*(*P*_1,1_) − *φ*(*P*_1,0_) are negative, and |*φ*(*P*_0,1_) − *φ*(*P*_0,0_)| *>* |*φ*(*P*_1,1_) − *φ*(*P*_1,0_)| (Figure 7D).

Therefore, in this example, 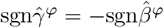.

Note, that if *φ* is increasing convex down, −*φ* is decreasing convex up; and if *φ* is increasing convex up, −*φ* is decreasing convex down. Change of the sign of the function inverts the directions of both 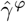 and 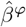 (see (15) and (16)). Thus, those pairs of transformations, induce the same relation between the signs of 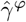 and 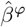.

Finally, change of the sign of 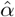, inverts the relation between the magnitudes of differences *φ*(*P*_0,1_) − *φ*(*P*_0,0_) and *φ*(*P*_1,1_) − *φ*(*P*_1,0_), which, for a given transformation, results in an inverted relation between the signs of 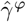 and 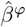. We summarize all possible cases in Table 1.

## Appendix B

Consider the following model of phenotype *Y*:

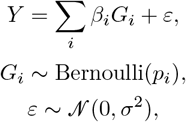

where *G*_*i*_ indicates the presence of an alternative allele at variant *i, β*_*i*_ is the effect of this allele on *Y*, and *ε* is the environmental noise. We assume that the genotypes are independent: ∀_*i*≠*j*_*G*_*i*_ ╨ *G*_*j*_, and that the environmental noise is homoskedastic: ∀_*i*_Var[*ε*|*G*_*i*_ = 0] = Var[*ε*|*G*_*i*_ = 1]. Note that, unlike in Appendix A where no specific generating model is assumed, here we assume this is the actual generating process. Without loss of generality, let *σ*^2^ = 1.

Suppose that if the level of *Y* is high, treatment *E* is administered:

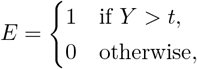

where *t* is some threshold. When applied, treatment *E* changes the level of *Y* by *α*:

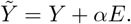

Suppose further that we observe phenotype 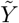 and test the effect 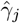 of the interaction between variant *G*_*j*_ and environmental factor *E* on this phenotype:

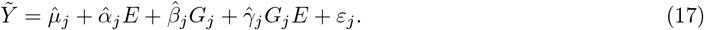

We prove that the sign of 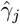 is determined by the sign of the main effect 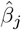 of *G*_*j*_ (Claim 2).

Note that coefficients 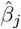 and 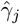 can be related to empirical conditional expectations of 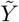:

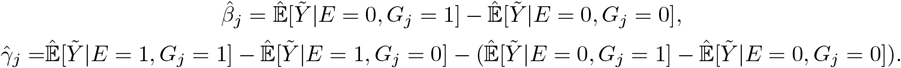

Furthermore, note that phenotype 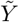 conditioned on the value of *E* has a truncated normal distribution, and its conditional mean is given by:

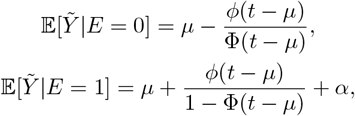

where *ϕ* is the probability density function and Φ is the cumulative distribution function of the standard normal distribution, and *µ* is the mean of *Y*. Furthermore, the value of *µ* depends on the genotype *G*_*j*_:

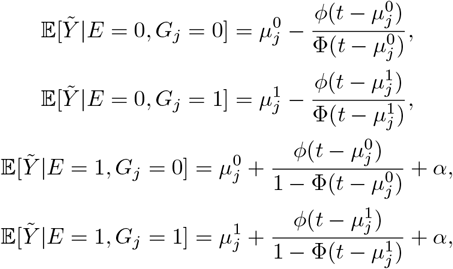

Where 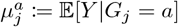.

To simplify the above expressions, we denote the inverse Mill’s Ratio *λ*(*x*) ≔ *ϕ*(*x*)*/*Φ(*x*), and note that *ϕ*(*x*)*/*(1 − Φ(*x*)) = *ϕ*(−*x*)*/*Φ(−*x*) = *λ*(−*x*), because *ϕ* is even, and Φ(−*x*) = 1 − Φ(*x*). Furthermore, we define points 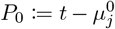 and 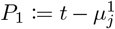, and express the estimated effects 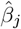 and 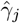 in (17) as a function of these points:

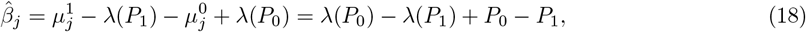

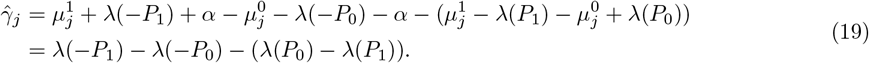

The function *λ* is decreasing and strictly convex down [22] (Figure 8). As a result, the order of points *P*_0_ and *P*_1_ determines the signs of 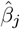 and 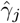. Without loss of generality, let *t >* 0. Since *t* distinguishes ”high” from ”normal” levels of phenotype *Y*, it is reasonable to assume that means 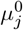 and 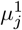 are smaller than *t* (note that variants *G*_*i*_ are independent); that is, any individual SNP does not result in high enough *Y* to receive the treatment, as it is likely for any polygenic trait. There are therefore two possible cases:

1. 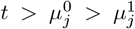, which imposes the following order on the points used in definitions (18) and (19): −*P*_1_ *<* −*P*_0_ *< P*_0_ *< P*_1_ (Figure 8A). Given this order and the properties of *λ*, we have: 1) *P*_0_ − *P*_1_ *<* 0, *λ*(*P*_0_) − *λ*(*P*_1_) *>* 0, and |*λ*(*P*_0_) − *λ*(*P*_1_)| *<* |*P*_0_ − *P*_1_|, which implies that 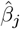 is negative; and 2) 0 *< λ*(*P*_0_) − *λ*(*P*_1_) *< λ*(−*P*_1_) − *λ*(−*P*_0_), which implies that 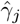 is positive.
2. 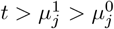, which results in: −*P*_0_ *<* −*P*_1_ *< P*_1_ *< P*_0_ (Figure 8B). Given this order and the properties of *λ*, we have: 1) *P*_0_ − *P*_1_ *>* 0, *λ*(*P*_0_) − *λ*(*P*_1_) *<* 0, and |*λ*(*P*_0_) − *λ*(*P*_1_)| *<* |*P*_0_ − *P*_1_|, which implies that 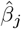 is positive; 2) *λ*(−*P*_1_) − *λ*(−*P*_0_) *< λ*(*P*_0_) − *λ*(*P*_1_) *<* 0, which implies that 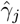 is negative.

**Figure 8:**
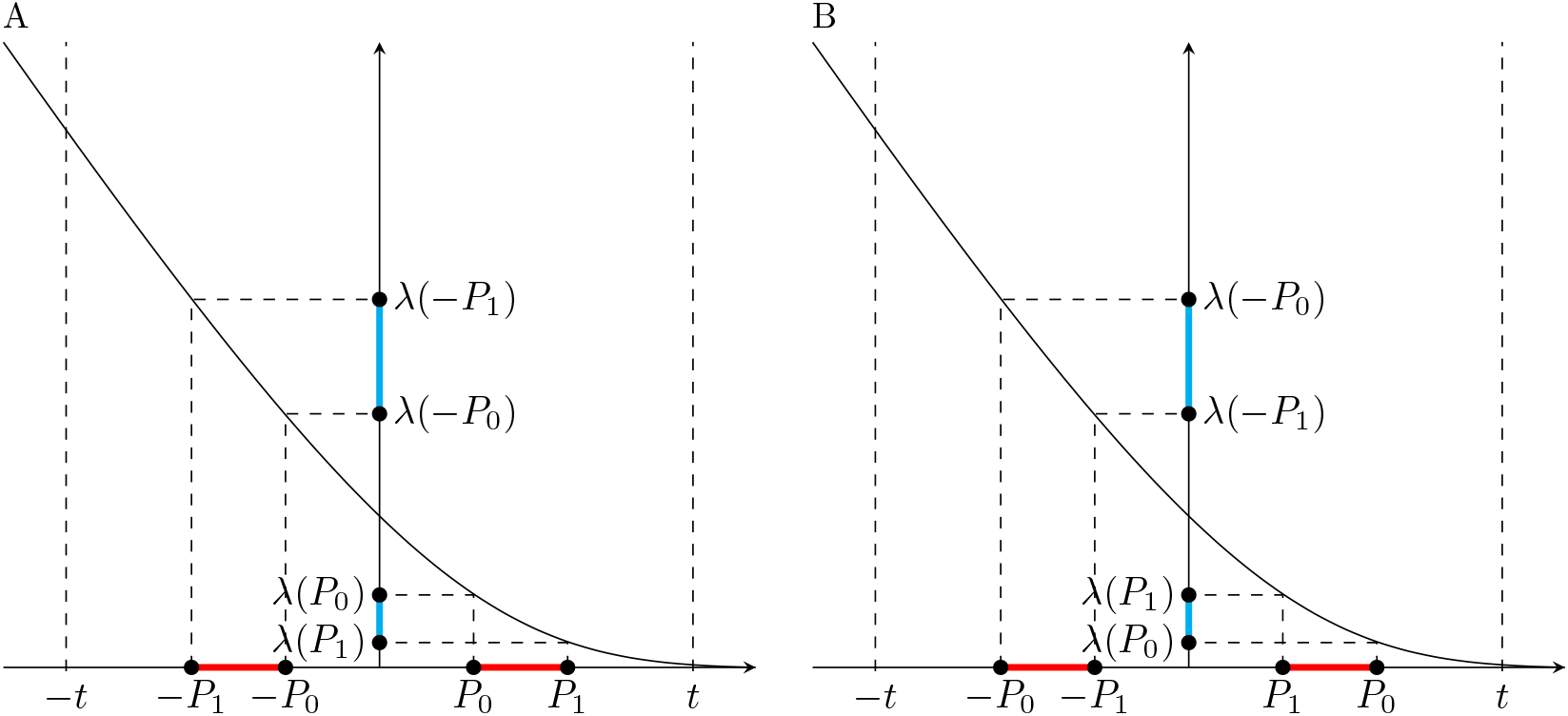
Endogenous treatment effects induce sign-consistent *G* × *E* effects. (A) The x-axis shows the quantities 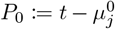 and 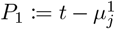 defined for phenotype *Y* affected by the haploid genetic variant *G*_*j*_, where 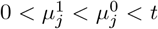. The main effect of *G*_*j*_ and the effect of *G*_*j*_*E* on phenotype *Y*, after treatment *E* is applied to reduce levels of *Y* that exceed threshold *t*, can be expressed as functions of *P*_0_ and *P*_1_ and their images under the inverse Mill’s Ratio *λ*(*P*_0_) and *λ*(*P*_1_). The signs of those functions are dependent. (B) Similar to (A), but when 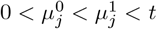.

We have, therefore, shown that:

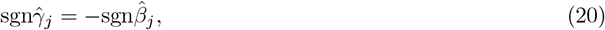

which means that the estimated effects 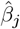 and 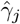 in regression (17) have opposite directions. It can be analogously shown that relation (20) holds when treatment *E* is administered whenever the level of phenotype *Y* is below threshold *t*.

## 8 Acknowledgments

This research was possible through funding from the National Institutes of Health. This research used the UK Biobank Resource under application 33127. We thank the participants of the UK Biobank for making this work possible.

## Supplementary Material

### The logistic transformation

Consider the logistic function:

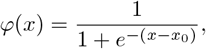

and four points *P*_0,0_, *P*_0,1_, *P*_1,0_, and *P*_1,1_, such that *P*_0,1_ − *P*_0,0_ = *P*_1,1_ − *P*_1,0_ = *d* (Figure 9A).

**Figure 9:**
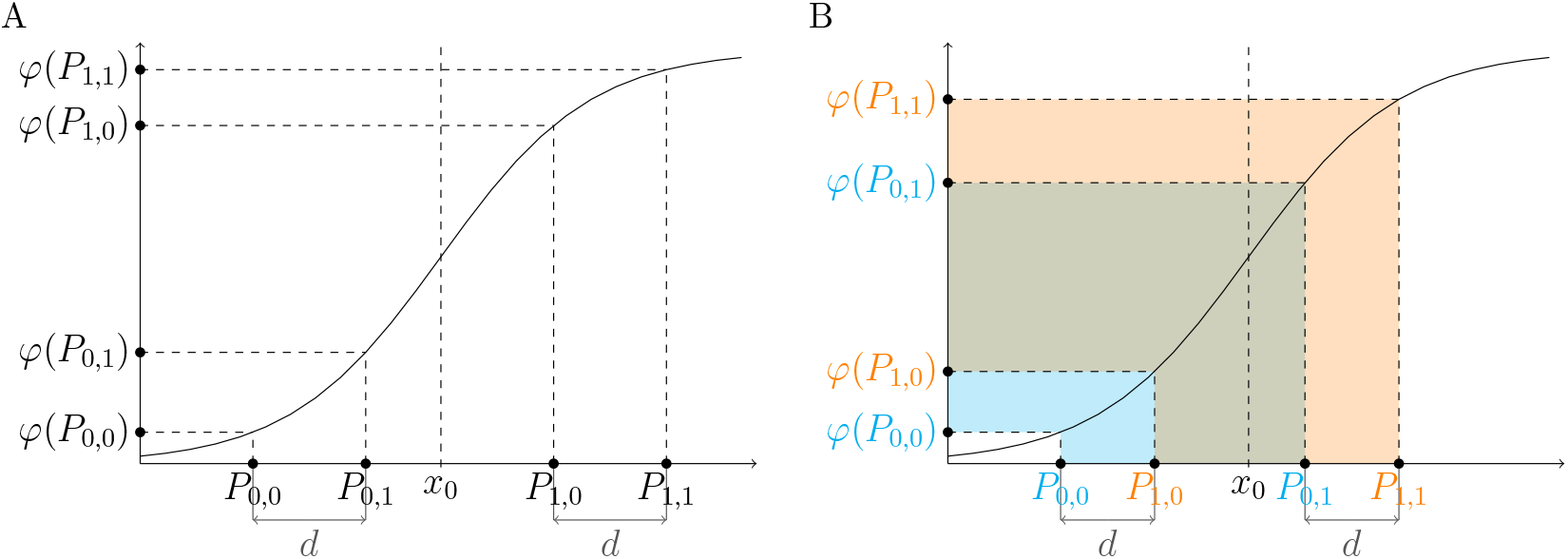
The effect of the values *P*_0,0_, *P*_0,1_, *P*_1,0_, and *P*_1,1_ on the sign of expression: *φ*(*P*_1,1_) − *φ*(*P*_1,0_) − (*φ*(*P*_0,1_) − *φ*(*P*_0,0_)), where *φ* is the logistic function. (A) When the order of the input values is *P*_0,0_ *< P*_0,1_ *< x*_0_ *< P*_1,0_ *< P*_1,1_. (B) When the order of the input values is *P*_0,0_ *< P*_1,0_ *< x*_0_ *< P*_0,1_ *< P*_1,1_.

We examine the sign of 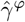, the value defined below, as a function of the values of points *P*_0,0_, *P*_0,1_, *P*_1,0_, and *P*_1,1_:

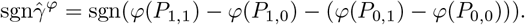

There are six possible cases (without loss of generality, we assume that *P*_0,1_ − *P*_0,0_ *>* 0 and *P*_1,0_ − *P*_0,0_ *>* 0):

1. If all the points *P*_*a,b*_ are smaller than *x*_0_, transforming them with *φ* is equivalent to transforming them with an increasing convex down function. As shown in Appendix A, this results in 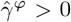 (Table 1 and Figure 2A).
2. If all the points *P*_*a,b*_ are larger than *x*_0_, transforming them with *φ* is equivalent to transforming them with an increasing convex up function. As shown in Appendix A, this results in 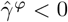 (Table 1 and Figure 2C).
3. If *P*_0,0_ and *P*_0,1_ are smaller than *x*_0_, but *P*_1,0_ and *P*_1,1_ are larger than *x*_0_, the sign of 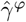 depends on the relation between distances |*P*_0,1_ − *x*_0_| and |*P*_1,0_ − *x*_0_| (Figure 4A). Note that *φ*^*′*^(*x*_0_ + *x*) = *φ*^*′*^(*x*_0_ − *x*), and that the derivative of *φ* decreases as the distance from *x*_0_ increases. Recall also our assumption that *P*_0,1_ − *P*_0,0_ = *P*_1,1_ − *P*_1,0_. Therefore:
  - If |*P*_0,1_ − *x*_0_ | *>* |*P*_1,0_ − *x*_0_ |, then 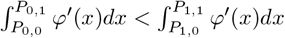. Consequently, 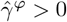.
  - If |*P*_0,1_ − *x*_0_ | *<* |*P*_1,0_ − *x*_0_ |, then 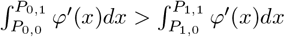. Consequently, 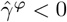.
4. Similarly as in point 3, if *P*_0,0_ and *P*_1,0_ are smaller than *x*_0_, but *P*_0,1_ and *P*_1,1_ are larger than *x*_0_, the sign of 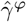 depends on the relation between distances |*P*_0,1_ − *x*_0_| and |*P*_1,0_ − *x*_0_| (Figure 9B). Note that:

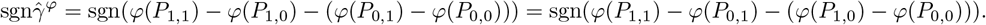

We use the same argument as in point 3, but this time the differences we compare are *P*_1,1_ − *P*_0,1_ and *P*_1,0_ − *P*_0,0_, instead of *P*_1,1_ − *P*_1,0_ and *P*_0,1_ − *P*_0,0_. This comparison yields:
  - If |*P*_0,1_ − *x*_0_| *>* |*P*_1,0_ − *x*_0_|, then 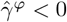.
  - If |*P*_0,1_ − *x*_0_| *<* |*P*_1,0_ − *x*_0_|, then 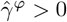.
5. If *P*_0,0_ is smaller than *x*_0_, and *P*_0,1_, *P*_1,0_ and *P*_1,1_ are larger than *x*_0_, then—since *φ*^*′*^ decreases as the distance from *x*_0_ increases—*φ*(*P*_0,1_) − *φ*(*P*_0,0_) is greater than *φ*(*P*_1,1_) − *φ*(*P*_1,0_), and consequently 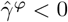.
6. Conversely to point 5, when only *P*_1,1_ is larger than *x*_0_, and *P*_0,0_, *P*_0,1_ and *P*_1,0_ are smaller than *x*_0_, *φ*(*P*_0,1_) − *φ*(*P*_0,0_) is smaller than *φ*(*P*_1,1_) − *φ*(*P*_1,0_), and consequently 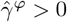.

### TxEWAS in the UK Biobank

The TxEWAS presented in this work were performed following the Sadowski et al. protocol [23]. The studied UK Biobank population of 342,257 unrelated white British individuals was identified by performing the steps described by Sadowski et al. [6]

We imputed gene expression into the UK Biobank using eQTL weights trained in 48 tissues of The Genotype-Tissue Expression (GTEx v7) project, linked by the TxEWAS protocol [23]. Hierarchical FDR (hFDR *<* 10%) was used to account for multiple hypothesis testing across genes and tissues [31, 32].

Individuals who took statins were identified by codes: 1140861958, 1140861970, 1141146138, 1140888594, 1140888648, 1140910632, 1140910654, 1141146234, 1141192410, 1141192414, 1141188146, 1140881748, and 1140864592 in the UK Biobank field 20003-0.0-47. Smoking status was derived from the UK Biobank field 20116-0.0 by encoding the ”current” category as 1, and the categories of ”never” and ”previous” as 0.

For all tested outcomes except testosterone, we discarded measurements greater than five standard deviations from the mean, with the assumption that such extreme levels were results of non-modeled circumstances. The distribution of testosterone levels was bimodal, but the sign-consistency pattern for this phenotype presented in Figure 5A remained similar after inverse normally transforming it.

We included age, sex, birth date, Townsend deprivation index, and the first 16 genetic PCs [25] as covariates in our studies. All non-binary covariates were standardized (transformed to mean zero, variance one) before calculating interaction variables.

### Heteroskedasticity

Heterogeneity of phenotype variance across values of genetic and/or environmental factors (heteroskedasticity) can produce false *G* × *E* [33]. Since large differences of phenotypic variance between environments are common, conditional heteroskedasticity is an important source of statistical artifacts in *G* × *E* studies [6, 29]. It is therefore important that interaction models account for it.

In the presence of environment-conditional heteroscedasticity, testing multiple genetic variants for interaction with the environmental factor in a simple linear regression model results in an inflated or deflated false positive rate (FPR), depending on the relation between group size and phenotypic variation [29]. We demonstrate this in simulation and derive a formula revealing this relation in a univariate model.

Consider a binary environmental variable that divides observations into two groups of sizes *n*_0_ and *n*_1_, and phenotype variances 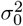 and 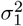. We simulated data for 10,000 such observations. The phenotype was drawn from Gaussian distributions with mean zero and corresponding group variances. We independently drew 200 genotypes from a binomial distribution *B*(2, *p*) with *p* representing the minor allele frequency, which was drawn uniformly from a range between 0.1 and 0.5. For each genotype, two *G* × *E* models were fitted: 1) a simple linear regression model that included the genotype, the environmental factor and the product of those two as covariates (OLS), and 2) a double generalized model (DGLM) that used the same covariates to model the mean effects, and the environmental variable to model the variance effects.

We ran this simulation for selected values of the ratio 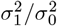, and calculated the FPR for the *G* × *E* effect as the proportion of simulations where the nominal p-value for this effect was less than 0.05 (Figure 10A). The simulation shows that if the smaller (larger) group is characterized by the larger (smaller) variance of the response, the FPR for the OLS is inflated (deflated). Within a realistic range of parameter values, the FPR can reach zero or increase threefold. On the other hand, if the groups have equal sizes, the model is well calibrated. Figure 10B shows p-value distributions for one such simulation with realistic variance ratios 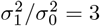 and 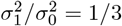[6].

**Figure 10:**
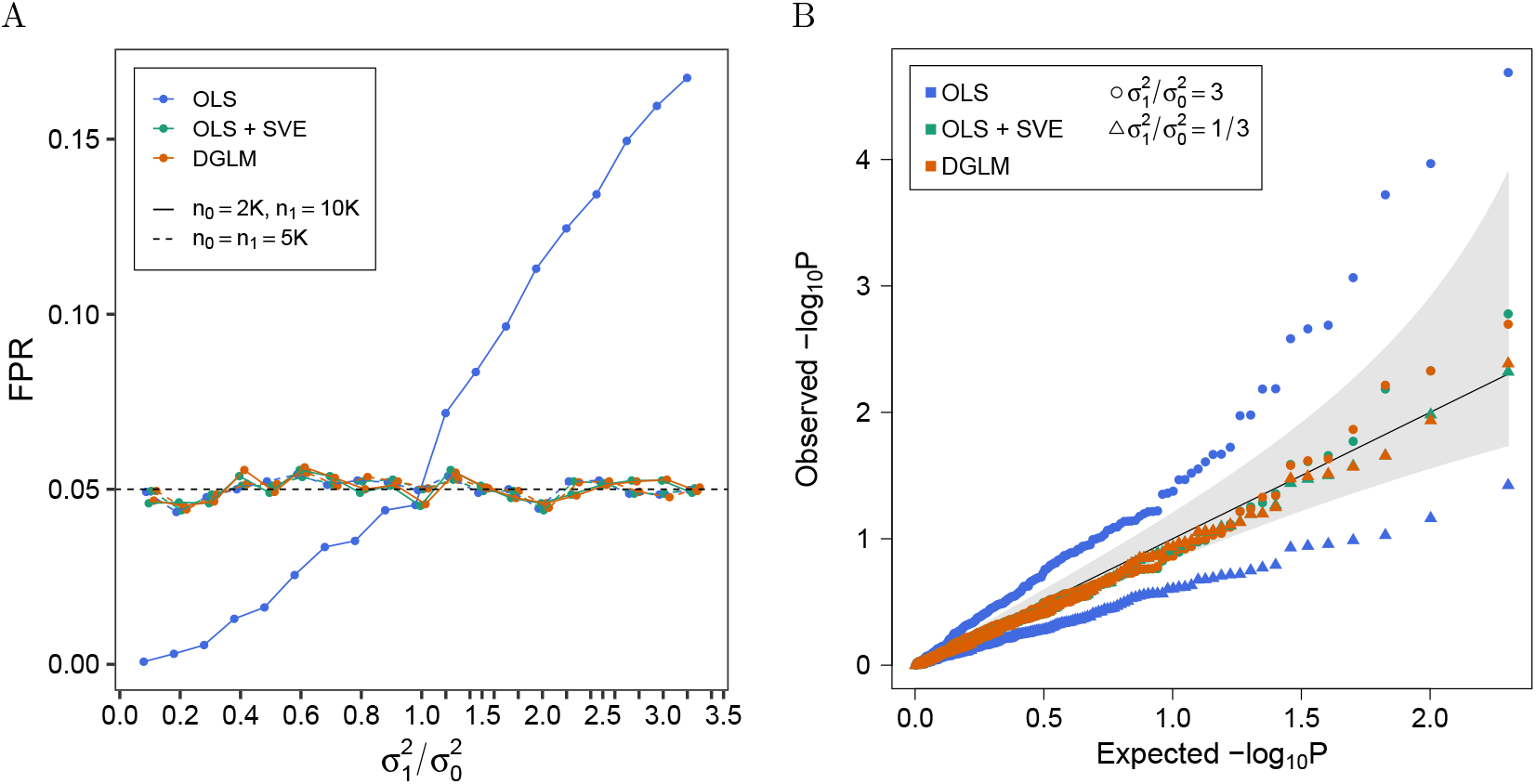
Evaluation of the type I error rate for the *G* × *E* effect estimated with the OLS model, the OLS model using robust standard errors (OLS + SVE) and the DGLM. (A) False positive rate (FPR) of *G* × *E* as a function of the ratio between phenotype variances in two environments: unexposed (of size *n*_0_ and phenotype variance 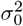), and exposed (of size *n*_1_ and phenotype variance 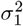). The nominal FPR of 5% is marked by the black dashed line. (B) Quantile-quantile plot comparing the null expected p-values (x-axis) to the observed *G* × *E* p-values when 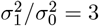 (circles) and when 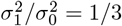 (triangles).

Below is an analytical demonstration of how this bias arises. Consider the following linear regression model:

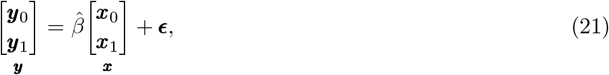

where:

- ***y*** is a size *N* vector generated by concatenating vectors ***y***_0_ and ***y***_1_, and centering. Individually, ***y***_0_ (***y***_1_) is a vector of size *n*_0_ (*n*_1_), whose elements were independently drawn from a Gaussian distribution with mean zero and variance 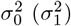.
- ***x*** is a size *N* vector generated by concatenating a size *n*_0_ vector of zeros (***x***_0_) and a size *n*_1_ vector of ones (***x***_1_), and centering. We use *k*_0_ ≔ −*n*_1_*/N* and *k*_1_ ≔ 1 − *n*_1_*/N* = *n*_0_*/N* to refer to the values of ***x***_0_ and ***x***_1_, respectively, after centering.
- ***ϵ*** is a size *N* vector of residuals.

The variance of the estimate 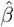 is given as:

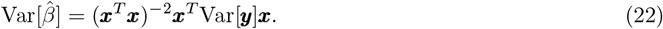

Given the assumptions of model (21), it can be expressed as:

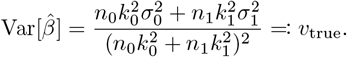

Recall that *k*_0_ = −*n*_1_*/N*, and *k*_1_ = *n*_0_*/N*. We can make those substitutions to simplify the above expression to:

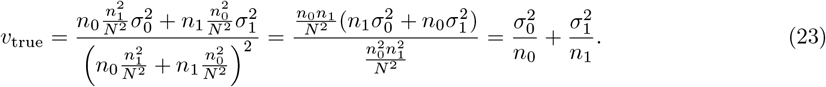

However, the simple linear regression (SLR) model assumes homoskedasticity (Var[***y***] = *σ*^2^***I***, where ***I*** is an identity matrix), in which case (22) simplifies to:

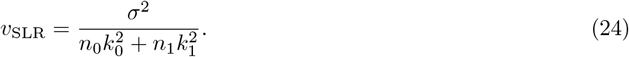

The parameter *σ*^2^ is then estimated as:

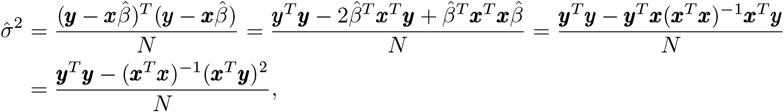

where we use the fact that 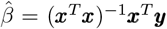. Given the assumptions of model (21), this can be computed as:

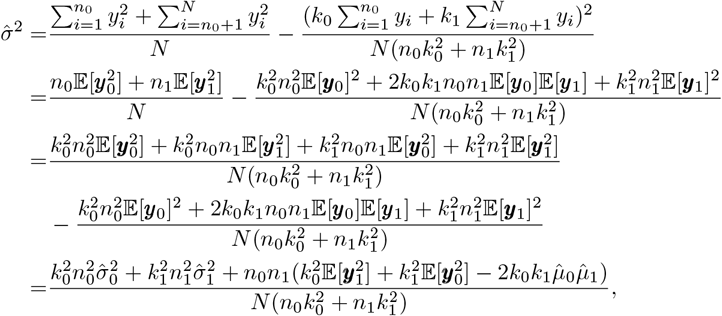

Where 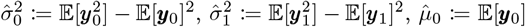, and 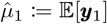. This expression is simplified when substituting *k*_0_ for −*n*_1_*/N*, and *k*_1_ for *n*_0_*/N* :

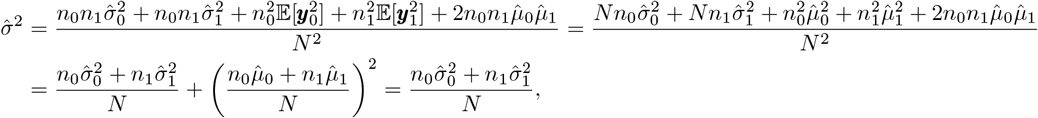

where the last equality follows from the fact that ***y*** is centered. Therefore, following (24), the SLR estimates the variance of 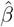 as:

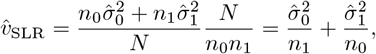

which is different from the estimate of the true variance, 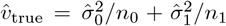, as derived in (23). In particular, examination of the difference between these estimates:

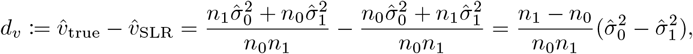

shows that:

- if *n*_0_ *> n*_1_ and 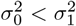, then *d*_*v*_ *>* 0, which means that the *t*-statistic computed with SLR is inflated;
- if *n*_0_ *> n*_1_ and 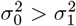, then *d*_*v*_ *<* 0, which means that the *t*-statistic computed with SLR is deflated;
- if *n*_0_ = *n*_1_, then 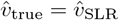, which means that the *t*-statistic computed with SLR is well-calibrated.

The described bias can be removed by incorporating robust standard errors estimated with the sandwich variance estimator (SVE) into the OLS (Figure 10). Alternatively, the data can be correctly modeled with the DGLM.

